# Repair of DNA double-strand breaks after low radiation doses in childhood cancer survivors and matched cancer-free individuals

**DOI:** 10.64898/2025.12.22.695953

**Authors:** J Mirsch, RN Cordoni, C Schmitt, A Schulze, D Galetzka, S Zahnreich, P Scholz-Kreisel, T Hankeln, M Marron, M Blettner, CM Ronckers, H Schmidberger, M Löbrich

**Affiliations:** Radiation Biology and DNA Repair, Technical University of Darmstadt, Darmstadt, Germany; Institute of Medical Biostatistics, Epidemiology and Informatics, University Medical Centre of the Johannes Gutenberg University Mainz, Mainz, Germany; Department of Radiation Oncology and Radiation Therapy, University Medical Centre of the Johannes Gutenberg University Mainz, Mainz, Germany; Radiation Epidemiology and Radiation Risk, Federal Office for Radiation Protection, München (Neuherberg), Germany; Institute of Organismic and Molecular Evolution, Molecular Genetics and Genome Analysis, Johannes Gutenberg University Mainz, Mainz, Germany; Department of Epidemiological Methods and Etiological Research, Leibniz Institute for Prevention Research and Epidemiology - BIPS, Bremen, Germany; German Childhood Cancer Registry, Division of Childhood Cancer Epidemiology, Institute of Medical Biostatistics, Epidemiology and Informatics, University Medical Centre of the Johannes Gutenberg University Mainz, Mainz, Germany; Division of CAYA Cancer Survivorship Research; German Cancer Research Center. Heidelberg, Germany

**Keywords:** DNA double-strand breaks, low radiation doses, genetic predisposition, childhood cancer, radiation risk

## Abstract

DNA double-strand breaks (DSBs) which arise in G1- or G0-phase normal human cells are repaired by non-homologous end-joining (NHEJ), a pathway which is important for cell survival but can cause mutations at the break sites. DSB repair by NHEJ is very efficient at high damage levels of 1 or more DSBs per cell, much less efficient at lower damage levels and almost absent if only ∼0.05 DSBs per cell are induced. Repair at this low DSB level can be induced if cells are pre-treated prior to DSB induction with low concentrations of hydrogen peroxide, suggesting that the intracellular radical level modulates the efficiency of DSB repair at low damage levels. Here, we have investigated if the inefficiency of repair at low DSB levels contributes to the carcinogenic potential of DSBs or, counterintuitively, may serve as a mechanism to limit cancer development. We have analyzed the repair of high and low levels of radiation-induced DSBs in primary fibroblasts from 136 childhood cancer survivors, half of whom developed a second independent tumor later in life, and compared it to the response of primary fibroblasts from 68 individually matched cancer-free individuals. Although childhood cancer survivors and cancer-free individuals repaired DSBs at high damage levels equally efficiently, their response to low DSB levels differed drastically. While repair in cancer-free individuals was nearly absent at a level of ∼0.05 DSBs per cell, childhood cancer survivors repaired DSBs at this low damage level as efficiently as after high damage levels. These results suggest that most of the cancer survivors analyzed here carry a genetic predisposition that affects their response to low levels of DSBs. They also support the idea that the absence of repair observed in cancer-free individuals represents a mechanism limiting cancer formation.

## Introduction

DNA double-strand breaks (DSBs) are an important class of DNA lesions since unrepaired breaks lead to loss of genetic information and cell death. The repair of DSBs can restore the genetic information but can also cause mutations, which represent initiating events for cell transformation and cancer development [1–5]. The predominant pathway for DSB repair is non-homologous end-joining (NHEJ), particularly in G1- or G0-phase cells which represent the majority of the cells in a human body [3, 6]. NHEJ has no intrinsic feature to restore genetic information lost at the break site and hence can lead to small deletions that can alter gene function [3, 7]. Moreover, NHEJ can erroneously connect break ends from different DSBs which leads to genomic rearrangements, a hallmark of many cancer cells [8]. Thus, repairing DSBs by NHEJ is important to allow cell survival, but carries the risk of causing cancer.

Inherited defects in DNA repair genes often lead to clinical syndromes arising from excessive cell death but can also manifest as a predisposition of the individuals to cancer development [9–12]. Examples of inherited defects in DSB repair genes include individuals with a syndrome caused by mutations in the NHEJ factor DNA ligase 4 [13, 14]. This syndrome is associated with microcephaly due to excessive neuronal cell death, with immunodeficiency due to impaired development of B lymphocytes and with a predisposition to cancer development [10, 11, 13]. Cells from such individuals exhibit genomic instability and extreme sensitivity to DSB-inducing agents [13–15]. Other examples of inherited defects in DSB repair genes are heterozygous mutations in BRCA1 or 2 which are involved in homologous recombination, a second major DSB repair pathway [12, 16, 17]. Individuals who carry heterozygous BRCA1 or BRCA2 mutations are at a substantially higher risk to develop tumors (mainly breast and ovarian cancer) than non-carriers [18, 19], yet without extreme sensitivities to DSB-inducing agents. These cases show that a proper response to DSBs is key to warrant organismic development and limit cancer formation.

Most molecular and cellular studies to investigate DSB repair processes have been carried out using high doses of radiation or other DNA-damaging agents. However, humans are generally not exposed to high doses of DNA-damaging agents and only very few DSBs arise in any given cell under physiological conditions, at least during the G1- or G0-phase of the cell cycle when DSB repair occurs by NHEJ [3, 6]. Hence, it is important to understand how NHEJ repairs low levels of DSBs and if the repair efficiency differs between high and low damage levels. To address this question, we and others have previously irradiated fibroblasts in the G1 phase of the cell cycle with doses of 2.5 mGy - 2 Gy X-rays, inducing 0.0625 - 50 DSBs per cell [20–23]. DSB repair by NHEJ is very efficient at damage levels of 1 or more DSBs per cell but, unexpectedly, declines at lower damage levels, leading to a complete failure to repair DSBs after an X-ray dose inducing only ∼0.05 DSBs per cell [20, 22, 23].

Subsequent studies have revealed that the repair efficiency by NHEJ at low damage levels can be stimulated by pre-treating cells prior to irradiation with 10 µM hydrogen peroxide, a concentration which does not induce DSBs itself but substantially enhances the cellular reactive oxygen level [20]. This suggests that conditions which modulate the homeostatic redox system are critical for the efficiency of repair after physiologically relevant DSB levels. Lastly, the inefficient repair of low levels of DSBs is not only observed for cells in culture since mice irradiated with low radiation doses also fail to repair the induced DSBs [20, 24]. Collectively, these studies established that DSBs induced by low doses of DNA-damaging agents remain unrepaired for many days, raising the question about the physiological relevance of this phenomenon. In particular, it is unclear if the lack of repair at low DSB levels contributes to the carcinogenic potential of DSBs or if it represents a mechanism limiting cancer formation.

We have addressed this question in the current manuscript by analyzing the repair efficiency at low DSB levels in a large group of individuals who developed cancer during childhood and compared it to the DSB repair efficiency of matched individuals who never had cancer. The rationale for this approach is that childhood cancer represents a group of rare diseases, with only 175 new cases per 1 million children in Germany per year [25]. The most common types of childhood cancer are leukemia (mainly acute lymphoblastic leukemia), lymphoma (Hodgkin and non-Hodgkin lymphoma), CNS tumors, and a variety of solid tumors (among others, neuroblastoma, Wilms tumor and osteosarcomas). Despite the early onset of childhood cancer, only a small fraction of all cases (about 10%) has hitherto been associated with a specific genetic predisposition [9, 26–30]. However, because of the rarity of these diseases or reduced penetrance of disease-causing genes [28], a given genetic predisposition likely confers only a small increase in risk for a specific childhood cancer type and may, therefore, be virtually impossible to identify based on analysis of the family cancer history. Recent publications indicate that, as sequencing technologies advance, more genetic predispositions are likely to be identified in the upcoming years [9, 26, 27, 31, 32].

Here, we analyzed the repair of DSBs in fibroblasts from childhood cancer survivors and cancer-free individuals. We found that cancer-free individuals fail to repair low levels of DSBs while childhood cancer survivors repair DSBs at low damage levels almost as efficiently as after higher damage levels. This suggests that (i) childhood CNS tumors and lympho-hematopoietic cancers may be associated with a genetic predisposition affecting the repair of low DSB levels and that (ii) the lack of repair at low damage levels might represent a mechanism limiting cancer formation.

## Results

### Experimental design and automated foci scoring

To investigate the hypothesis that childhood cancer is associated with a genetic pre-disposition that manifests as an altered efficiency to repair low levels of DSBs, we analyzed fibroblasts collected as part of the KiKme-nested case-control study [33]. Adult individuals who survived a common childhood primary neoplasm (leukemia, lymphoma or CNS tumor) were identified from the German Childhood Cancer Registry and invited to participate in the study. Matched cancer-free individuals were also participating, providing a unique collective of carefully matched individuals. In total, we obtained fibroblasts from skin biopsies of 204 individuals, including 136 long-term childhood cancer survivors, 68 of whom developed a second primary neoplasm (SPN). The 68 individuals who developed a first primary neoplasm (FPN) but no SPN were individually matched to the 68 individuals with an SPN and further matched to 68 cancer-free individuals (0PN, see Fig. 1A). Additional information about the participants and the Kikme study is provided in table 1 and Marron *et al*. [33].

**Figure 1:**
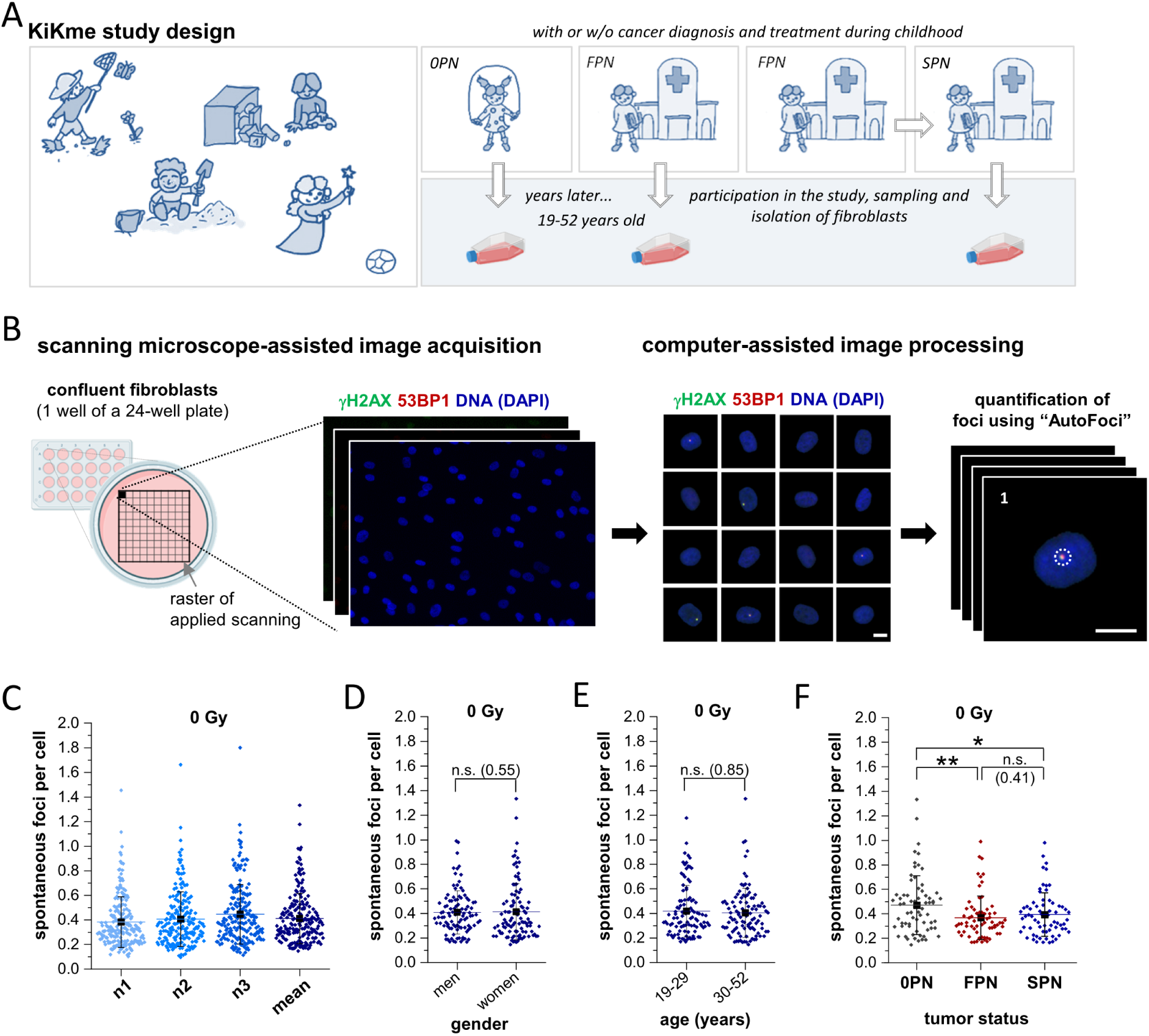
Experimental design, image processing and analysis of spontaneous foci in a large group of 204 individuals. (**A**) Graphical visualization of the study design. Individuals who survived childhood cancer were identified by the German Childhood Cancer Registry. 68 childhood cancer survivors with a primary neoplasm were carefully matched to 68 individuals who developed a second independent primary neoplasms after an FPN in childhood. Matching criteria included age, gender and year and tumor type at first diagnosis. 68 cancer-free individuals were matched to these pairs by age and gender, creating matched triplets. For the analysis of DSB repair, fibroblasts were isolated from skin samples of each participant. (**B**) Graphical visualization of the experimental design. Cells from 7 triplets were seeded randomly into one 24-well plate and analyzed simultaneously. Overview images were processed using our Cellect tool to generate 3000-5000 single cell images per sample and γH2AX/53BP1 foci were counted automatically using AutoFoci software. Throughout all experiments, samples were blinded and their group identification (0PN, FPN, SPN) and treatment condition (radiation dose) were unknown. The scale bars represent 10 µm. (**C**) Quantification of spontaneous foci in cells from untreated samples. Data for individual experiments (n1-n3) and the mean of the three experiments are shown. Each point represents one individual. 18 of the 612 data points (three experiments times 204 individuals) for individual experiments were excluded from the analysis due to insufficient staining quality. The population mean and s.d. are shown in black. (**D**-**F**) Analysis of spontaneous foci grouped according to sex (D), age at recruitment (E) or cancer history (F). Each data point represents the mean of the 3 independent experiments of an individual. The population mean and s.d. are shown in black. Statistical analysis was performed using a linear mixed model, and all statistical parameters are described in table 2A. The illustration of the cell culture flask (for panel A) and the 24-well plate (for panel B) were created using bioRENDER.com. 0PN, cancer-free individuals; FPN, first primary neoplasm; SPN, second primary neoplasm; *, p ≤ 0.05; **, p ≤ 0.01; n.s., not significant.

**Table 1:**
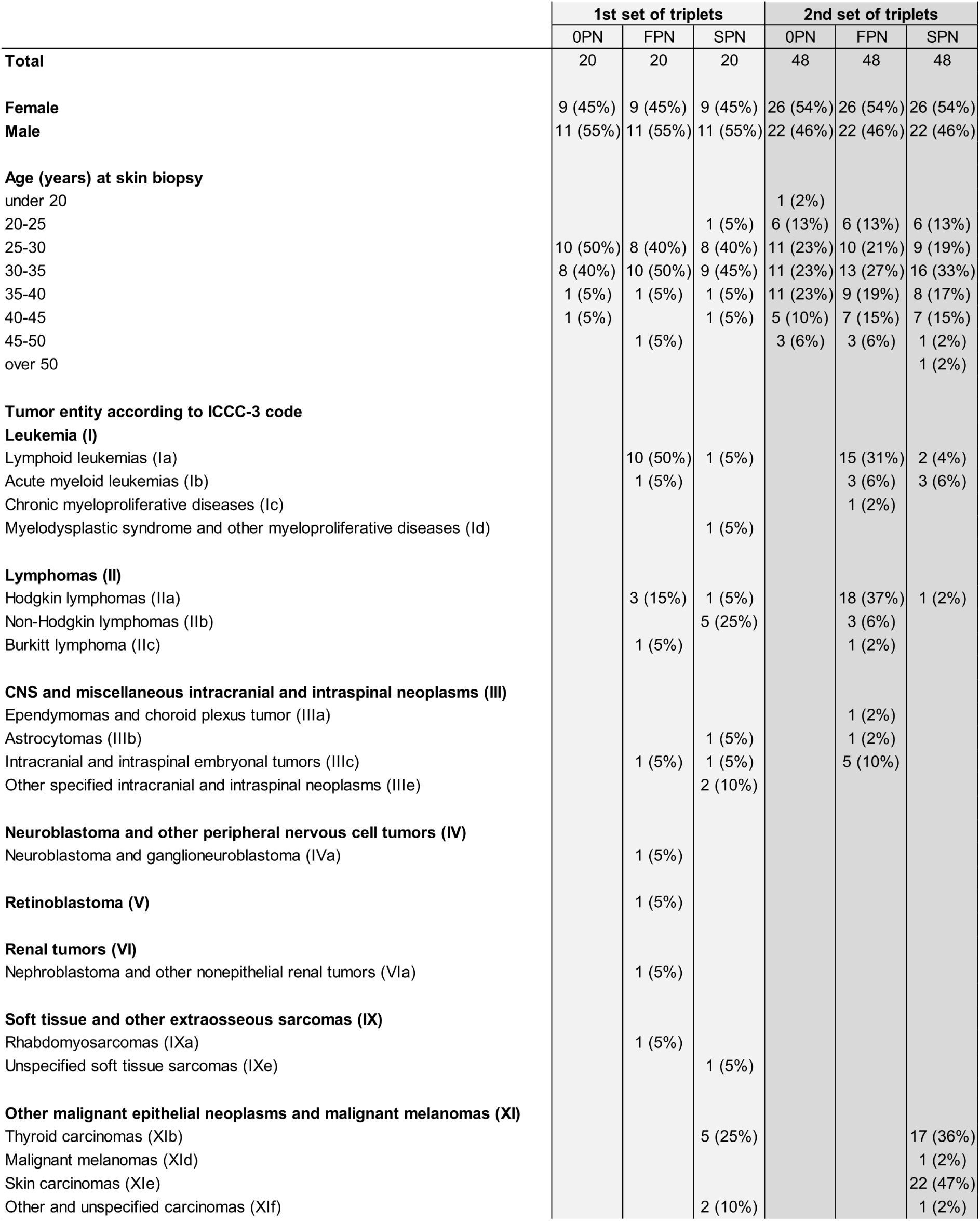
Overview of the characteristics of participants of the KiKme study. The KiKme study design including the recruitment process for childhood cancer survivors and cancer-free individuals, their matching and the sampling of punch skin biopsies to isolate the fibroblasts are described in Marron *et al.* [33]. Because of the matching criteria, the spectrum of first neoplasms of individuals with an SPN corresponds to that of individuals with an FPN. Note that the type of SPN is unknown in one individual of the main study and that four individuals developed a third independent neoplasm which is not included in this table. Gender, age at skin biopsy and tumor entity according to international classification of childhood cancer, third edition (ICCC-3)-code [57] are listed.

**Table 2:** Overview of statistical parameters for the analysis of spontaneous foci levels. (**A**) We first applied a linear mixed model (LMM) which revealed a significant effect for cancer history group but not for age and gender (p = 0.85 and p = 0.55, respectively). We then performed post hoc tests to assess all combinations of cancer history groups at 0 mGy. The results showed that spontaneous foci levels are significantly higher in cancer-free individuals than in childhood cancer survivors but not significantly different between the FPN and SPN groups. (**B**) Three separate linear models were applied to the spontaneous foci levels of individuals sorted by FPN type. Models for leukemia and lymphoma as FPN revealed a significant effect of FPN type but not of age or gender. However, the model for ‘CNS and others’ revealed no effect on FPN type and age, but on gender (p = 0.03). We then performed post hoc tests to assess differences in spontaneous foci numbers between childhood cancer survivors and their corresponding control groups. The results showed that spontaneous foci levels in cancer-free individuals are significantly higher than in childhood cancer survivors with Leukemia or Lymphoma but not with ‘CNS and others’. (**C**) We first applied an LMM to the datasets of spontaneous foci shown in Figs. S4B and C, which revealed a significant effect for cancer history group but not for age or gender. We then performed post hoc tests to assess differences in foci numbers between childhood cancer survivors and the control group. The results confirmed that spontaneous foci levels are significantly higher in cancer-free individuals than in childhood cancer survivors. Dose, cancer history group, effect estimate beta, s.e.m. and p value are listed. 0PN, cancer-free individuals; xPN, individuals surviving at least one primary neoplasm in childhood; FPN, first primary neoplasm; SPN, second primary neoplasm.

| A | dose | contrast |  | estimate | s.e.m. | p value |
| --- | --- | --- | --- | --- | --- | --- |
| 1 | 0 mGy | xPN | 0PN | -0.192 | 0.067 | 0.0046 |
| 2 | 0 mGy | FPN | 0PN | -0.225 | 0.078 | 0.0042 |
| 3 | 0 mGy | SPN | 0PN | -0.160 | 0.078 | 0.0404 |
| 4 | 0 mGy | SPN | FPN | 0.065 | 0.078 | 0.4055 |

| B | dose | contrast |  | estimate | s.e.m. | p value |
| --- | --- | --- | --- | --- | --- | --- |
| 1 | 0 mGy | xPN Leu | 0PN (Leu gr.) | -0.231 | 0.107 | 0.0336 |
| 2 | 0 mGy | xPN Lym | 0PN (Lym gr.) | -0.235 | 0.111 | 0.0379 |
| 3 | 0 mGy | xPN CNS | 0PN (CNS gr.) | -0.115 | 0.139 | 0.4130 |

| C | dose | contrast |  | estimate | s.e.m. | p value |
| --- | --- | --- | --- | --- | --- | --- |
| 1 | 0 mGy | xPN (S4B) | 0PN (S4B) | -0.218 | 0.060 | 0.0004 |
| 2 | 0 mGy | xPN (S4C) | 0PN (S4C) | -0.175 | 0.066 | 0.0093 |

To measure the repair of DSBs after low radiation doses in a large number of samples, we applied a previously described automated analysis system [21]. Fibroblasts were grown to a confluent monolayer to minimise the formation of spontaneous DSBs due to DNA replication. We used 24-well plates to simultaneously analyze samples from different individuals and performed three biological replicates. We irradiated the samples with X-ray doses from 2.5 to 100 mGy, fixed the cells 24 h after irradiation and stained them for γH2AX and 53BP1 which form co-localising foci at DSBs in G0- or G1-phase cells. We obtained 100 overview images from about 3-5 thousand cells per sample (Fig. 1B), segmented each overview image to generate single cell images and applied our program AutoFoci [21]. AutoFoci detects objects containing foci and non-specific background signals and records their object properties such as intensity, size and sharpness. A unique combination of these object properties shows a bimodal distribution that can be used to distinguish true foci from background signals.

### High inter-individual variations of spontaneous DSBs

First, we analyzed unirradiated cells and observed a wide range of spontaneous DSBs in the samples from different individuals. While numbers as low as 0.1 foci per cell were observed in some samples (Fig. 1C), samples from other individuals exhibited more than 1 focus per cell. 90% of the samples exhibited between 0.18 and 0.87 foci per cell and 50% of the samples between 0.27 and 0.5 foci per cell. Foci numbers for different individuals were consistent between the three independent experiments, suggesting that the inter-sample variations largely represent inter-individual variations rather than experimental uncertainties (Fig. S1A).

To substantiate this notion, we selected five individuals with considerably different numbers of spontaneous DSBs and analyzed three additional samples for each individual. The foci numbers obtained in these additional experiments were highly consistent with the results from the screen that involved all 204 individuals (Fig. S1B).

To investigate potential causes underlying the large inter-individual variations, we grouped the individuals either by sex, by the age at the time the biopsies were taken, or by whether they developed an SPN, an FPN or no cancer. The analysis of 99 men and 105 women showed no association between spontaneous foci numbers and sex (Fig. 1D). Also, two similar-sized age groups, 19-29 years (99 individuals) and 30-52 years (105 individuals), showed no significant difference in spontaneous foci numbers (Fig. 1E). However, and most interestingly, the 68 cancer-free individuals exhibited significantly more foci than the 68 FPN and 68 SPN survivors (Figs. 1F and S1C). This result is in line with the idea that individuals who develop childhood cancer might carry a genetic predisposition which affects their cellular response to low DSB levels.

### Assessing foci levels after low radiation doses

Next, we investigated the response of the 204 individuals to low doses of X-rays. We irradiated confluent fibroblasts and analyzed the foci values at 24 h after irradiation to assess the repair capacity of each individual. The irradiated samples showed a broad spectrum of foci, similar to that observed for unirradiated samples (Fig. 2A). Although not directly visible by eye, the average foci value of all irradiated samples was clearly elevated above the background level of spontaneous foci (Fig. 2A), indicating the presence of unrepaired DSBs at 24 h after irradiation. To better assess each person’s individual repair capacity, we subtracted the spontaneous foci value of each individual from the foci values of the samples after irradiation. This led to a much more confined distribution of unrepaired radiation-induced foci although some individuals exhibited numbers that were substantially higher than expected or even negative (Fig. 2B). We have previously established that X-rays induce foci in confluent fibroblasts linearly with dose at a rate of about 25 foci per Gy [20, 22, 34]. Here, we selected five individuals, measured foci numbers at 15 min after irradiation with doses between 2.5 and 100 mGy and confirmed the induction rate of 25 foci per Gy with little to no variation between the individuals (Figs. 2C and S2A). Based on this induction rate, we plotted the range for the expected unrepaired radiation-induced foci as a grey box into the measured data, with the upper end of the box showing the numbers if all radiation-induced foci would stay unrepaired and the lower end of the box showing the numbers if all foci would be repaired (Fig. 2B).

**Figure 2:**
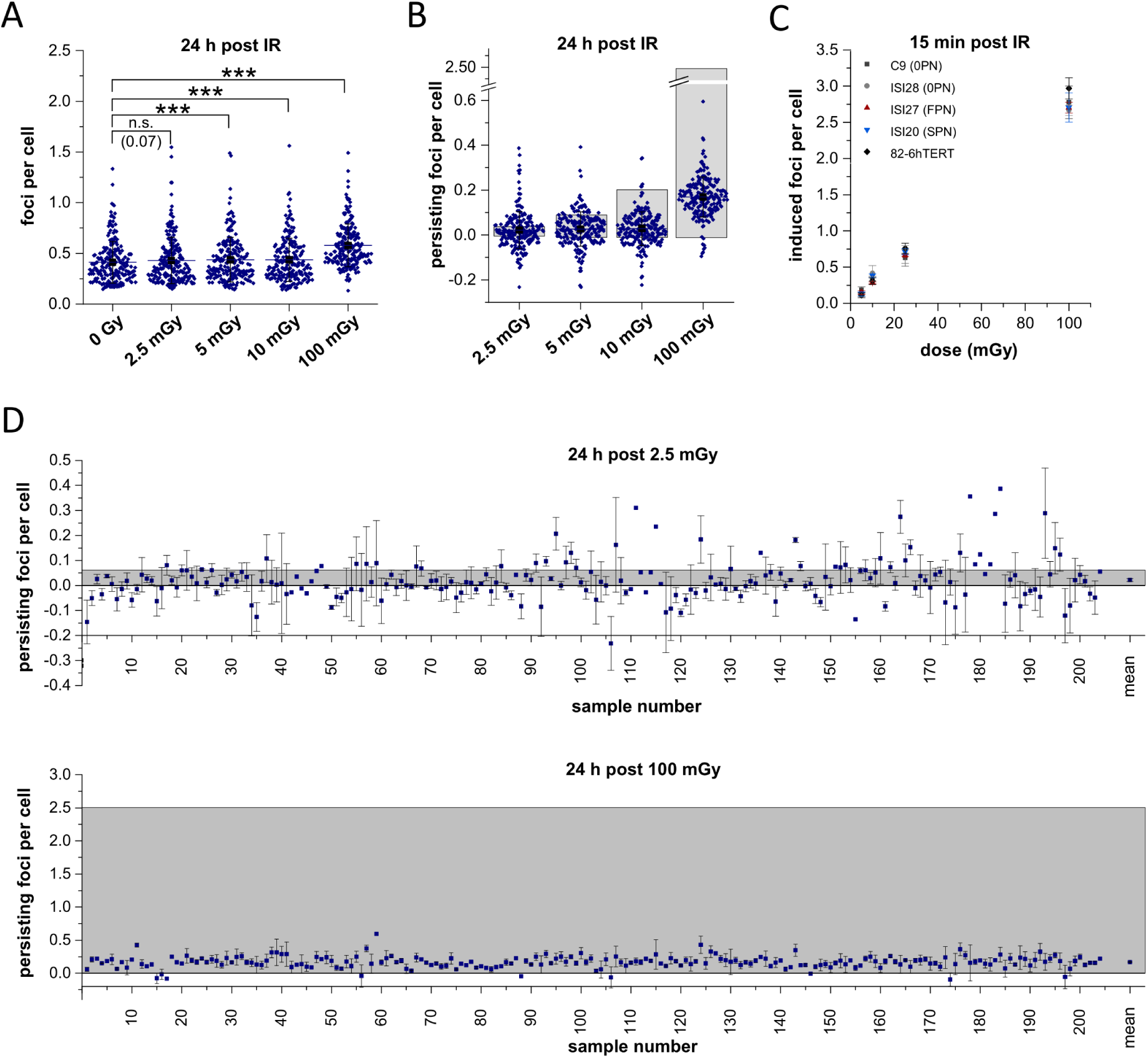
Foci levels after low radiation doses. (**A**) Quantification of foci in irradiated cells using AutoFoci. Data for the mean of the three experiments are shown. Each point represents one individual. 64 out of the 2448 irradiated samples (four doses times three experiments times 204 individuals) were excluded from the analysis due to an insufficient number of cells or poor staining quality. For the 18 out of the 612 unirradiated samples that were excluded (see Fig. 1C), the irradiated samples of the same experiment were also excluded. In total, 154 out of the 3060 samples were excluded, in which case the presented mean values are derived from only two (for 146 samples) or one (for 4 samples) independent experiments. For each irradiation condition, the population mean and s.d. are shown in black. The values for unirradiated cells are taken from figure 1C. (**B**) Persisting foci after irradiation calculated from the data in panel A. For this, the foci level of unirradiated samples was subtracted from the foci level after irradiation for each individual experiment and the data for the independent experiments were averaged for each individual. For each irradiation dose, the population mean and s.d. are shown in black. The grey-shaded box for each dose indicates a theoretical range of foci obtained by calculating the number of induced foci per cell for each dose based on an induction rate of 25 foci per Gy. (**C**) Foci induction after irradiation in cells from four selected individuals and an established cell line which differ substantially in their spontaneous foci levels (see Fig. S2A). The foci level of unirradiated samples was subtracted from the foci level at 15 min after irradiation for each individual experiment and the data for the independent experiments were averaged for each individual. The mean and s.d. are shown. A linear regression analysis of all data points provided the following parameters: f(x) = 0.027x (+/- 0.0005), R^2^=0.99. (**D**) Persisting foci after irradiation with 2.5 and 100 mGy. The data from panel B is replotted, showing the average number of persisting foci for the 204 individuals and the s.e.m. if three independent measurements are available. The population mean and s.e.m. are shown as the last point in each row. Corresponding plots for 5 and 10 mGy are shown in figure S2B. Statistical analysis was performed using a linear mixed model, and all statistical parameters are described in table 3. ***, p ≤ 0.001; n.s., not significant.

**Table 3:**
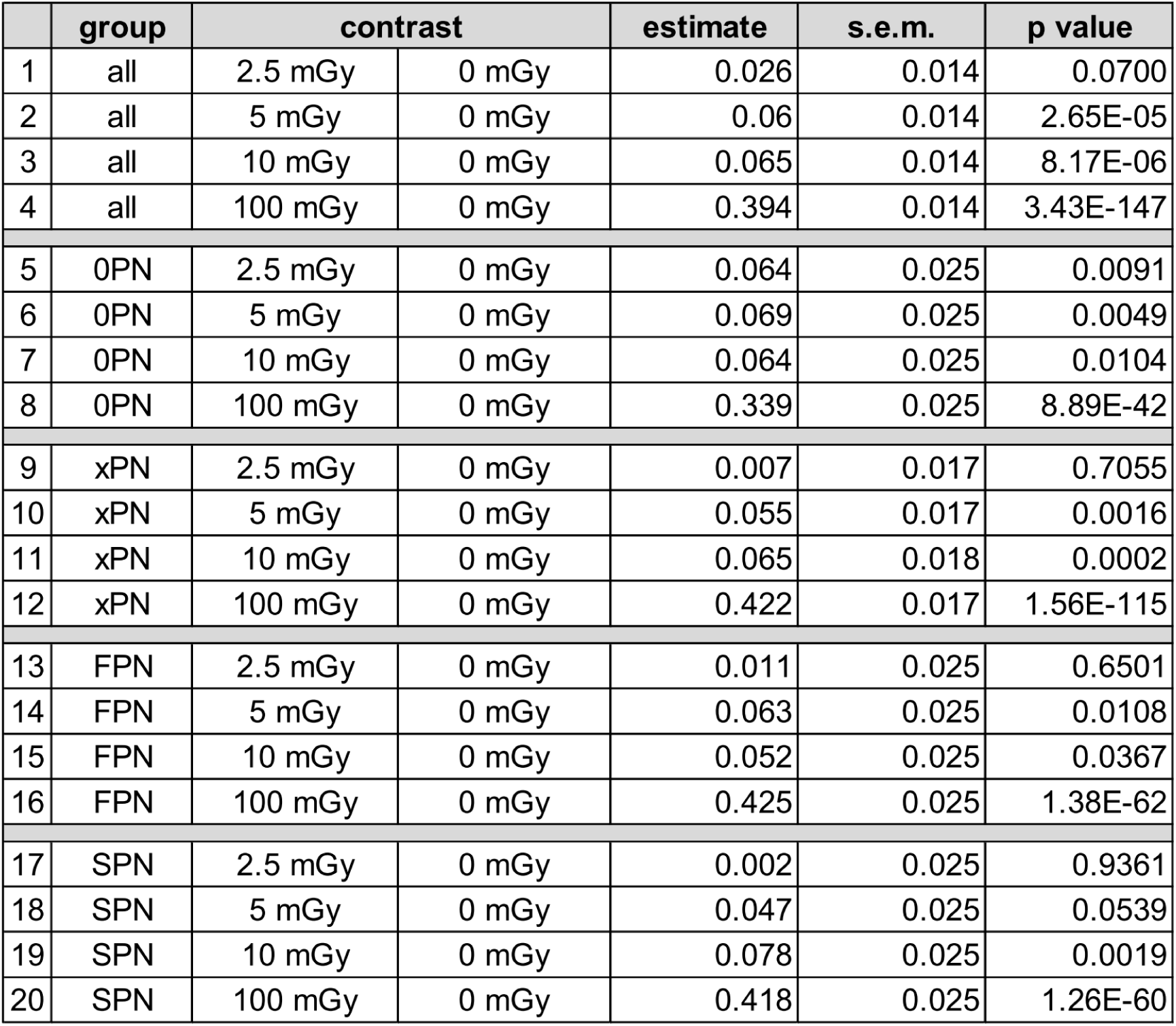
Overview of statistical parameters for the analysis of all foci levels. We first applied a linear mixed model which revealed a significant effect for cancer history group but not for age and gender (p = 0.88 and p = 0.63, respectively). We then performed post hoc tests to assess radiation-induced increases in foci levels. Foci levels of all individuals together increased significantly after doses of 5 mGy and higher and with a small p-value of 0.070 for 2.5 mGy (lanes 1-4). Foci levels of cancer-free individuals increased significantly for all doses (lanes 5-8). In contrast, foci levels of childhood cancer survivors increased significantly for doses of 5 mGy and higher (lanes 10-12) but not for a dose of 2.5 mGy (lane 9). Similar results were obtained when distinguishing between individuals affected by an FPN (lanes 13-16) or an SPN (lanes 17-20) among childhood cancer survivors. This analysis confirmed the lower number of persistent DSBs in childhood cancer survivors at 2.5 mGy. Dose, cancer history group, effect estimate, s.e.m. and p value are listed. 0PN, cancer-free individuals; xPN, individuals surviving at least one primary neoplasm in childhood; FPN, first primary neoplasm; SPN, second primary neoplasm.

We then replotted the data in Fig. 2B showing for each individual the number of unrepaired radiation-induced foci and the standard error derived from the three independent measurements (Figs. 2D and S2B). The measured foci numbers fell within the range of expected foci numbers for 35, 55, 65 and 98% of the individuals after doses of 2.5, 5, 10 and 100 mGy, respectively. The range of the measured foci numbers defined by the standard error overlapped with the range of expected foci in 74, 86, 85 and 100% of the individuals after doses of 2.5, 5, 10 and 100 mGy, respectively. This analysis showed that, at least for the three lowest doses, the uncertainty of the measured foci number for every one individual is too large to allow a comparison between single individuals. We, therefore, calculated for each dose the average foci number of all individuals and plotted them together with the standard errors derived from the differences between the 204 individuals in the same figures (Figs. 2D and S2B). These four average foci numbers all fell within the box of expected foci numbers, and their standard error was considerably smaller than the range of expected foci. We conclude that the experimental design with three independent foci measurements per individual is insufficient to accurately assess the radiation response of single individuals after low radiation doses but are capable to determine the average response of a group of individuals. Therefore, subsequent analyses compare average foci levels for groups of individuals.

### Childhood cancer survivors repair low dose damage better than tumor-free individuals

We first assessed the average repair efficiency of all individuals by dividing the number of unrepaired radiation-induced foci by the number of foci induced at the corresponding doses. We observed that 7% of the foci induced by 100 mGy remain unrepaired for 24 h, suggesting efficient DSB repair at this dose (Fig. 3A). Strikingly, the fraction of unrepaired foci increased to 11% after 10 mGy, 21% after 5 mGy and 36% after 2.5 mGy, confirming our previous observation that the efficiency for repairing radiation-induced DSBs decreases with decreasing doses [20–22]. We conclude that our automated high-throughput foci scoring system analyzing millions of cells from 204 different individuals recapitulates the finding that DSBs induced by low radiation doses are inefficiently repaired.

**Figure 3:**
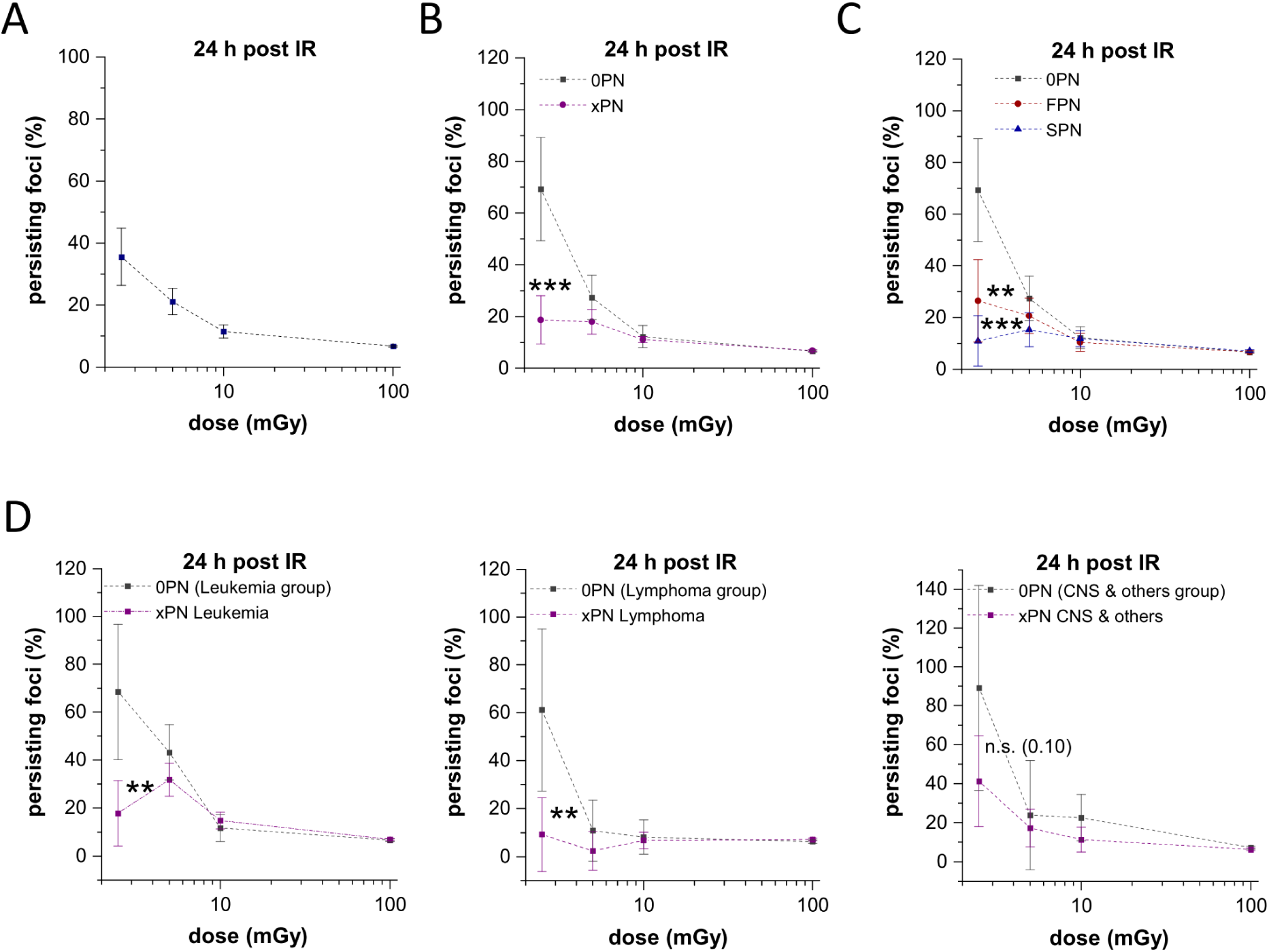
Childhood cancer survivors repair low levels of DSBs better than cancer-free individuals. (**A**) Fraction of persisting foci as a function of dose. The number of persisting foci per cell was divided by the expected number of induced foci per cell based on an induction rate of 25 foci per Gy. The population mean and s.e.m. are shown. (**B**, **C**) Fraction of persisting foci as a function of dose separated for cancer history: 68 cancer-free individuals and 136 childhood cancer survivors (B) or separated in 68 individuals with an FPN and 68 with an SPN (C). The population mean and s.e.m. are shown. (**D**) Fraction of persisting foci as a function of dose for 60 individuals with leukemia (left), 52 individuals with lymphoma (middle) and 24 individuals with ‘CNS and other tumor entities’ (right) as the first primary neoplasm together with the matched 30 (left), 26 (middle) and 12 (right) cancer-free individuals. The population mean and s.e.m. are shown. Statistical analysis was performed using a linear mixed model, and all statistical parameters are described in tables 4A (for panels B and C) and 4B (for panel D). 0PN, cancer-free individuals; xPN, individuals surviving at least one primary neoplasm in childhood; FPN, first primary neoplasm; SPN, second primary neoplasm; **, p ≤ 0.01; ***, p ≤ 0.001.

We next investigated if the observed impaired repair efficiency at low radiation doses differs between different groups of individuals and, similar to the analysis of the spontaneous foci levels, grouped the individuals by sex, by the age at the time the biopsies were taken, or by childhood cancer history. The analysis showed no effect of sex or age (Figs. S3A and B) but, strikingly, revealed that childhood cancer survivors repair DSBs induced by low radiation doses significantly better than their matched cancer-free individuals (Figs. 3B). Moreover, individuals affected by an SPN repaired slightly, but not statistically significantly, better than individuals affected by an FPN (Fig. 3C). Finally, we grouped all 136 childhood cancer survivors according to the type of the first neoplasm: leukemia (60 individuals), lymphoma (52 individuals) and ‘CNS tumors and others’ (24 individuals). Individuals affected by leukemia or lymphoma as FPN, but not CNS, repair low dose DSBs significantly better than their matched cancer-free controls (Figs. 3D and S3C).

To validate this finding, we first confirmed the automated foci scoring results by a manual evaluation of the microscopic images. We selected three SPN and three 0PN individuals who differed both in spontaneous foci number and repair efficiency and scored the foci by eye. The analysis of a total of ∼18.000 cells (6 individuals, 3 repeat experiments, 1000 cells per sample) confirmed the results from the automated scoring system (Fig. S4A). Next, we performed a statistical sensitivity analysis of the data. We removed either the 5 highest and the 5 lowest values per dose or identified outliers using the 3x interquartile range and removed those from the analysis. In both cases, the difference in spontaneous foci number and repair efficiency between childhood cancer survivors and cancer-free individuals remained statistically significant (Figs. S4B and C). This suggests that the observed effect cannot be explained by the response of a few individuals but rather indicates that the majority of childhood cancer survivors repair low dose DSBs better than the majority of cancer-free individuals.

## Discussion

We have designed this study to explore the physiological relevance of the phenomenon that DSB repair is very inefficient at damage levels substantially below 1 DSB per cell. Specifically, we asked if the near absence of repair at this damage level contributes to cancer development or whether it might represent a mechanism limiting cancer formation. To answer this question, we have invited adult childhood cancer survivors and matched cancer-free individuals to donate skin biopsies, established fibroblast cultures from the biopsies [33] and studied the response of these cells to low levels of X-ray-induced DSBs. We used X-rays to generate DSBs because ionizing radiation induces breaks from clusters of radicals which most closely resemble spontaneous DSBs arising from reactive oxygen species. In total, we measured the efficiency of DSB repair in fibroblast cultures from 204 individuals in three independent experiments. We assessed the DSB levels by the immunofluorescence analysis of γH2AX and 53BP1 foci and adapted an automated foci scoring system [21]. We performed all studies in a blinded manner and disclosed the identity of the samples to the experimenters only after all foci data were obtained.

The results from the analysis of 68 cancer-free individuals revealed about 10% unrepaired DSBs after doses of 10 and 100 mGy, about 30% unrepaired DSBs after 5 mGy and about 70% unrepaired DSBs after 2.5 mGy. This confirms our original finding obtained with a few selected fibroblast lines [20, 22] and establishes that the inefficient repair at low DSB levels represents a phenomenon of cancer-free individuals. In strong contrast, childhood cancer survivors exhibited less than 20% unrepaired DSBs at all analyzed doses, indicating efficient repair at all damage levels. A higher DSB repair efficiency after 2.5 mGy was also observed when the three groups of individuals with leukemia, lymphoma or ‘CNS tumors and others’ were compared separately to their corresponding control groups (0PN). Moreover, the DSB repair efficiency after 2.5 mGy was higher among individuals with SPNs (∼10% unrepaired DSBs) vs. individuals with FPNs (30% unrepaired DSBs). In all, childhood cancer survivors exhibited fewer spontaneous DSBs than cancer-free individuals, which is consistent with their higher efficiency of DSB repair after low doses. Collectively, this analysis supports the hypothesis that the development of childhood cancer is associated with an increased efficiency to repair low levels of DSBs.

We selected childhood cancer survivors for our study since we reasoned that tumors which develop at an early age are more likely to be associated with a genetic predisposition than tumors which form late during life. We included individuals who developed one of the three most common tumor types during childhood (leukemia, lymphoma or CNS tumor) and certain types of SPNs later in life (mean age at 2^nd^ diagnosis: 23 years). In particular, known therapy-related SPNs were included (thyroid cancer, leukemia and skin cancer). Although a high proportion of these specific second neoplasms likely arose from DNA damage inflicted by treatment of the first neoplasm, e.g. by chemotherapy, stem cell transplantation, or radiotherapy [35–39], we nevertheless expected that individuals who developed two independent tumors early in life might be even more likely to carry a genetic predisposition than individuals who developed only one neoplasm. The observation that individuals with SPNs repair low DSB levels better than individuals with FPNs is consistent with this idea. Thus, the result of our study suggests that the enhanced DSB repair efficiency of childhood cancer survivors at low DSB levels is caused by a certain form of genetic predisposition.

How can a genetic predisposition enhance the repair efficiency at low DSB levels? It is important to consider in this context that both groups, childhood cancer survivors as well as cancer-free individuals, efficiently repair DSBs by NHEJ at higher DSB levels (after 10 and 100 mGy). Also previous work with a sub-set of this case-control study did not report differences in survival, formation of chromosomal aberrations or gene expression changes after high radiation doses between childhood cancer survivors and cancer-free individuals [40, 41]. This suggests that the predisposition is unlikely to affect the main components of the DSB repair machinery, since this would be expected to change the repair efficiency at all damage levels. Instead, we propose that the genetic predispositions underlying childhood cancer formation specifically affect cellular systems or pathways which activate the DSB repair machinery in situations where it is inactive in normal individuals. Since we have previously observed that pre-treating cells with hydrogen peroxide, at a concentration which does not induce DSBs by itself but substantially enhances the intracellular radical level, activates DSB repair after low radiation doses [20], we speculate that alterations in the cellular redox system might underlie the predisposition of childhood cancer development.

The notion that an increased efficiency to repair DSBs at low damage levels is associated with cancer development might seem surprising. However, although DSB repair is important for cell survival, it can cause mutations and change gene function, especially if repair occurs by NHEJ, which has no intrinsic feature to restore sequence information lost at the break site [3, 7]. Thus, the decision to repair a break or not is a trade-off between warranting cell survival and potentially causing tumor-initiating mutations. After high damage levels, when the majority of the cells of an organism carry one or more DSBs, repair is necessary for the survival of an organism. But at low damage levels, when only a few cells have a break while the majority is undamaged, it might not be the best option for an organism to repair DSBs and risk the formation of mutations. Instead, the safer option could be to remove the cells with unrepaired DSBs and replace them with the division of undamaged cells. Such a concept is consistent with our observation that DSBs remain unrepaired in cells from cancer-free individuals when only 1 out of 20 cells carries a break but do get repaired when the level of DSBs increases. This concept is also consistent with our finding that childhood cancer is associated with a loss of this protective mechanism such that DSBs are repaired even if they arise in only a small fraction of all cells. The notion that cells with unrepaired DSBs might be replaced in normal individuals by the division of undamaged cells is supported by our earlier observation that cells with unrepaired DSBs disappear, while apoptotic cells and cells with micronuclei arise when cells re-enter the cell cycle [22].

The finding that the inefficiency of DSB repair by NHEJ at low damage levels might be beneficial has important implications. In the context of ionizing radiation, it is well established from many epidemiological studies that doses of 50 mGy and higher represent a carcinogenic risk which increases linearly with dose [42–45]. A linear relationship between dose and carcinogenic risk is consistent with a model that mutations which lead to cancer increase proportionally with dose. Since DSBs arise linearly with dose, it follows that the probability for a DSB to cause a mutation must be independent of dose. Our observation that the DSB repair efficiency of cancer-free individuals differs at low doses (<10 mGy) is inconsistent with this line of reasoning. Instead, our finding supports a model in which the mutation risk at low doses is smaller than what would be expected from a linear extrapolation at higher doses. Although carcinogenesis represents a multi-stage process involving several mutations and factors affecting tumor promotion (including mutations arising from processes other than the erroneous repair of DSBs), our findings appear to be inconsistent with a linear relationship between low dose radiation and carcinogenesis.

It is important to consider that we analyzed fibroblasts from childhood leukemia, lymphoma, and brain tumor survivors who underwent multi-modal tumor therapy. Many chemotherapeutic agents are known to induce oxidative stress or epigenetic changes which could potentially affect the efficiency of DSB repair [46–52]. Although we consider it unlikely that long-term cancer survivors exhibit persistent alterations in their metabolic system which affect the DSB repair efficiency, we cannot exclude the possibility that our observations were skewed by treatment-related effects. Moreover, since our analysis was limited to survivors of childhood cancer who reached adulthood, we equally cannot exclude the possibility of a “healthy survivor bias”.

## Materials and Methods

### Selection of study participants and isolation of primary fibroblasts

The population-based nested case-control study KiKme (German: “Krebserkrankungen im Kindesalter und molekulare Epidemiologie”; English: “Cancer in childhood and molecular-epidemiology”) was designed to investigate potential causes of childhood cancer, including genetic predispositions. The study was, therefore, performed with normal untransformed fibroblast samples and not with tumor cells, since these undergo various genetic changes during cancer progression. The study design, including the recruitment process of the cancer survivors and cancer-free individuals, their matching and the sampling of punch skin biopsies to isolate the fibroblasts is described in Marron *et al.* [33].

The first set of triplet samples included 40 participants with childhood cancer at any site (Table 1). For the second set, only participants with a first diagnosis of one of the three most common childhood cancers (leukemia, lymphoma and CNS tumors) were recruited. The second primary neoplasms were restricted to tumors which potentially could have been induced by radiation (thyroid cancer, skin cancer, including malignant melanoma or leukemia; Table 1). In addition to these and other regulatory requirements, participants were eligible if they were at least 18 years old and had survived at least 1 year after cancer diagnosis. First, individuals registered in the German Childhood Cancer Registry of the University Medical Centre of the Johannes Gutenberg University Mainz, Germany, with an SPN were identified and recruited (four individuals had developed 3 primary neoplasms). Individuals with an FPN were then matched to available SPN cases by age at recruitment (±5 years), sex, cancer site (ICCC-3), year of diagnosis (maximum range of ±7 calendar years), and age at diagnosis (maximum age range of ±4 years) using a risk set sampling approach. The mean age at 1^st^ diagnosis was 7 years (range 0-14 years) and at 2^nd^ diagnosis 23 years (range 5-46 years).

Additional cancer-free individuals for each matched pair of cancer survivors were recruited in the Department of Trauma Surgery of the University Medical Centre of the Johannes Gutenberg University Mainz, Germany. They were matched by age at recruitment (±5 years) and sex. Individuals with severe diseases were excluded from the study. For the first set of triplets, cancer-free individuals were recruited substantially later than the cancer survivors (>10 years) while for the second set, cancer survivors and cancer-free individuals were recruited at the same time (±2 years).

Skin samples of all childhood cancer survivors were taken at the Department of Radiation Oncology and Radiation Therapy of the University Medical Centre of the Johannes Gutenberg University Mainz, or their local dermatologists by punch biopsy under local anesthesia with a diameter of 3 mm at the cubital region. For cancer-free participants, the skin samples were taken at the University Medical Centre of the Johannes Gutenberg University Mainz during surgery in the scar region of a former small injury. The processing of the skin samples and the culture medium used are described in Marron *et al.* [33]. Fibroblasts were obtained and grown for 2-4 weeks before cryopreservation.

### Cultivation of primary human fibroblasts

The experimenters received the cryopreserved fibroblast samples in triplets for the repair study but were unaware of their cancer history until the end of the analysis. Fibroblasts were cultured in DMEM (low glucose, supplemented with 15% FCS, 1% NEAA, 100 units/ml penicillin, 0.1 mg/ml streptomycin, and amphotericin) at 37°C and 5% CO_2_, and cells were sub-cultured once a week at a ratio of 1:3 to 1:5 depending on the individual growth rate.

For each set of experiments, cells from 7 out of the 68 triplets were selected. Freshly thawed cells from these individuals were cultured for 2 weeks and used in the experiments at passages 9 to 11. For each individual, approximately 10,000 cells per cm^2^ were seeded into one well of 24-well glass bottom plates. Cells were grown for two weeks to obtain a non-dividing confluent cell layer. The cell culture medium was changed once during this time. The position of the samples on the plate was changed for each experiment so that potential differences due to the positioning were minimized.

### Irradiation of cells

Irradiation was performed with an X-ray machine (Titan Isovolt 160; General Electric) at 90 kV and 2 mA with a dose rate of 0.8-0.9 mGy/s (for doses up to 10 mGy) or 19 mA with a dose rate of 8-9 mGy/s (for doses between 10 and 100 mGy). The dose rate was determined and controlled with a dosimeter (DIADOS T60004; PTW Freiburg GmbH). Since cells were grown and irradiated on a glass surface, a dose correction factor of 3 was applied, which was determined experimentally according to a previous study [53].

### Fixation and immunofluorescence staining of γH2AX and 53BP1

Samples were analyzed 24 h after irradiation. Cells were washed with PBS and immediately fixed with 4% formaldehyde for 10 min. Cells were washed again with PBS and permeabilized with 0.2% TritonX100 in PBS. After three washing steps with the blocking solution Roti^®^Block (Carl Roth), cells were incubated for 30 min with Roti^®^Block at room temperature (RT).

For immunofluorescence staining of 53BP1 and γH2AX, cells were incubated with anti-53BP1 antibody (mouse; Upstate 24568) at 1:2000 and anti-γH2AX antibody (rabbit; Abcam GR1377) at 1:1000 in Roti^®^Block at 4°C overnight. After incubation with primary antibodies, cells were washed three times with Roti^®^Block and incubated with goat-anti-rabbit and goat-anti-mouse antibodies (Alexa Fluor 488 and 594, respectively; Invitrogen) at 1:500 or 1:1000 in Roti^®^Block for 1 h at RT in the dark. All subsequent steps were also performed in the dark. Cells were washed three times with Roti^®^Block and their DNA was stained with DAPI (0.4 µg/ml). Finally, cells were washed with ultrapure water and covered with ibidi mounting medium and a coverslip. The stained samples were stored at 4°C in the dark.

### Image acquisition and processing for computer-assisted foci scoring

All steps performed for image acquisition, image processing and computer-assisted DSB scoring using our program “AutoFoci” are described in Lengert *et al.* [21]. Briefly, a scan raster of 10x10 fields, each with a size of 1360x1024 pixels (439.28 x 330.75 µm^2^), was applied at a wide field microscope (Observer D1, Zeiss) with a 20x Plan-Apochromat objective (Zeiss). One field of view typically contained 50 to 100 cells. The image acquisition of the individual fields was automated utilizing the autofocus feature of the µManager software (Vale lab, UCSF). For each field, 1 image for DAPI and z-stacks of 5 images for each damage marker with a z-distance of 1.2 µm were captured. Using the customized ImageJ (NIH) tool Cellect, images were processed to generate single cell images of G1 cells while remaining S and G2 cells, as well as any potentially dying cells, were excluded by their abnormal nuclear morphology (DAPI) and high, pan-nuclear γH2AX signals.

For computer-assisted foci scoring, AutoFoci detects all objects, including foci as well as unspecific background signals, by local intensity maxima and additional object properties such as size and sharpness. These parameters were combined into a so-called object evaluation parameter (OEP). Using an inverse logarithmic graphic display of the OEP, the OEP shows a bi-modal object distribution with two maxima and a minimum in between that matches the transition between unspecific signals and foci. Since the distributions for background signals and foci overlap around the minimum, we implemented a short manual evaluation step to adjust the automatically estimated threshold. This brief manual intervention has the additional advantage that it allows a visual assessment of image and staining quality, preventing the usage of data with insufficient quality. Samples were excluded not only on the basis of poor staining but also if more than 50% of the cells did not meet the requirements set by AutoFoci, such as the signal-to-background ratio within the cells. During the manual adjustment step, the sample identity was unknown.

To validate the accuracy of the AutoFoci-assisted foci scoring data, manual foci counting was performed on a subset of samples (Fig. S4A). In this case, the co-localizing γH2AX and 53BP1 signals in G1 cells were counted as foci in a blinded manner using the overview images obtained with the 20x objective. Foci were counted in 1000 cells per sample.

### Foci induction by manual scoring of foci

To measure the induction of DSBs, selected samples were analyzed manually at 15 min after irradiation. The experiments were performed identically to the DSB repair measurements except that cells were grown on glass coverslips instead of 24 well plates. To identify G1 cells, EdU was added 1 h before irradiation and co-stained with DAPI, γH2AX and 53BP1 using a click-it reaction according to the manufacturer’s instructions (BaseClick). Since cells were growing on a coverslip, the coverslip was covered with mounting medium, placed on a slide and sealed with nail polish.

Manual foci counting was performed in a blinded manner at a fluorescence microscope from Zeiss using a 100x immersion objective (for Figs. 2C and S2A). Co-localizing γH2AX and 53BP1 signals in G1 cells were counted as foci. Foci were counted until a total number of 40 foci was reached and at least 40 cells were analyzed. Non-G1 cells with a positive EdU signal or high DAPI content were excluded from the analysis [21, 54].

DSB induction was also measured in 82-6hTert fibroblasts [54] which were cultured in MEM (supplemented with 15% FCS, 1% NEAA, 100 units/ml penicillin, 0.1 mg/ml streptomycin) at 37°C and 5% CO2, and sub-cultured once a week at a ratio of 1:10. The experiments were performed as described above, with cells grown for only one week before irradiation.

### Statistical analysis

The statistical analysis was performed in R (version 4.4.2.; R Core Team [55]). using the stats package for the linear models and the lmerTest package (version 3.1-3; Kuznetsova *et al*. [56]) for linear mixed models.

For the analysis of spontaneous foci levels (Tables 2A and C), the model was applied to the log-transformed number of foci at 0 mGy including fixed effects of group, as well as the covariates age and gender. As we have three measurements per individual and all measurements within an experimental group were carried out simultaneously, the individuals and their measurements within an experimental group were nested as random effects. For the analysis of foci levels after irradiation (Table 3), the model was applied to the log-transformed number of foci including fixed effects of group and dose and their interaction, as well as the covariates age and gender. The same model was applied to analyze the fraction of persisting foci (Tables 4A and C), which was calculated by dividing the difference in foci numbers between irradiated and unirradiated samples by the number of induced foci (25 per Gy).

The model was adjusted for the analysis of foci levels grouped by FPN type (leukemia, lymphoma and ‘CNS and others’), with the cancer-free individuals separated into three groups according to their matching. As this reduced the size of the overall dataset, the mean foci value for each individual was used instead of the three experimental measures per dose to reduce variability. For the analysis of spontaneous foci (Table 2B), the complexity was reduced to a linear model including the covariates age and gender, which was then applied to the log-transformed mean foci values. For the analysis of the fraction of persisting foci after irradiation (Table 4B), an LMM was applied to the mean foci value. Therefore, only the individuals were nested as random effects and fixed effects of group and dose, as well as the covariates age and gender, were included.

**Table 4:** Overview of statistical parameters for the analysis of persisting foci levels after irradiation. (**A**) We first applied a linear mixed model (LMM) which revealed a significant effect for cancer history group but not for age or gender (p = 0.78 and p = 0.68, respectively). We then performed post hoc tests to assess all combinations of cancer history groups for all doses. Cancer-free individuals showed foci levels at 2.5 mGy which are significantly higher than those of childhood cancer survivors (lanes 1, 5, 9) but not at higher doses (lanes 2-4, 6-8, 10-12). (**B**) Three separate LMMs were applied to the levels of persisting foci after irradiation for individuals sorted by FPN type. Models for leukemia and lymphoma as FPN revealed a significant effect of FPN type but not of age or gender. However, the model for ‘CNS and others’ revealed no effect on FPN type, age, or gender. We then performed post hoc tests to assess differences in the level of persisting foci between childhood cancer survivors and their corresponding control groups. Cancer-free individuals showed foci levels at 2.5 mGy which are significantly higher than those of childhood cancer survivors affected by Leukemia and Lymphoma (lanes 1, 5) but not at higher doses (lanes 2-4, 6-8). (**C**) We first applied an LMM to each dataset of persisting foci levels shown in Figs. S4B and C which revealed a significant effect for cancer history group but not for age or gender. We then performed post hoc tests to assess differences in the level of persisting foci between childhood cancer survivors and the control group. The results confirmed that cancer-free individuals exhibit foci levels at 2.5 mGy which are significantly higher than those of childhood cancer survivors. Dose, cancer history group, effect estimate, s.e.m. and p value are listed. 0PN, cancer-free individuals; xPN, individuals surviving at least one primary neoplasm in childhood; FPN, first primary neoplasm; SPN, second primary neoplasm.

| A | dose | contrast |  | estimate | s.e.m. | p value |
| --- | --- | --- | --- | --- | --- | --- |
| 1 | 2.5 mGy | xPN | OPN | -44.611 | 10.289 | 1.59E-05 |
| 2 | 5 mGy | xPN | OPN | -5.809 | 10.25 | 0.5710 |
| 3 | 10 mGy | xPN | OPN | -0.729 | 10.46 | 0.9445 |
| 4 | 100 mGy | xPN | OPN | 0.407 | 10.306 | 0.9685 |
| 5 | 2.5 mGy | FPN | OPN | -38.217 | 11.891 | 0.0013 |
| 6 | 5 mGy | FPN | OPN | -2.984 | 11.827 | 0.8009 |
| 7 | 10 mGy | FPN | OPN | -1.314 | 12.098 | 0.9136 |
| 8 | 100 mGy | FPN | OPN | 0.526 | 11.917 | 0.9648 |
| 9 | 2.5 mGy | SPN | OPN | -51.005 | 11.862 | 1.87E-05 |
| 10 | 5 mGy | SPN | OPN | -8.634 | 11.838 | 0.4659 |
| 11 | 10 mGy | SPN | OPN | -0.144 | 12.119 | 0.9905 |
| 12 | 100 mGy | SPN | OPN | 0.287 | 11.925 | 0.9808 |
| 13 | 2.5 mGy | SPN | FPN | -12.787 | 11.865 | 0.2814 |
| 14 | 5 mGy | SPN | FPN | -5.65 | 11.824 | 0.6328 |
| 15 | 10 mGy | SPN | FPN | 1.17 | 12.197 | 0.9236 |
| 16 | 100 mGy | SPN | FPN | -0.239 | 11.983 | 0.9841 |

| **B** | **dose** | **contrast** |  | **estimate** | **s.e.m.** | **p value** |
| --- | --- | --- | --- | --- | --- | --- |
| 1 | 2.5 mGy | xPN Leu | 0PN (Leu gr.) | -50.470 | 15.701 | 0.0014 |
| 2 | 5 mGy | xPN Leu | 0PN (Leu gr.) | -11.241 | 15.701 | 0.4746 |
| 3 | 10 mGy | xPN Leu | 0PN (Leu gr.) | 3.288 | 15.701 | 0.8343 |
| 4 | 100 mGy | xPN Leu | 0PN (Leu gr.) | 0.417 | 15.701 | 0.9788 |
| 5 | 2.5 mGy | xPN Lym | 0PN (Lym gr.) | -51.844 | 18.207 | 0.0048 |
| 6 | 5 mGy | xPN Lym | 0PN (Lym gr.) | -8.424 | 18.207 | 0.6440 |
| 7 | 10 mGy | xPN Lym | 0PN (Lym gr.) | -1.385 | 18.207 | 0.9394 |
| 8 | 100 mGy | xPN Lym | 0PN (Lym gr.) | 0.872 | 18.207 | 0.9618 |
| 9 | 2.5 mGy | xPN CNS | 0PN (CNS gr.) | -47.538 | 28.759 | 0.1014 |
| 10 | 5 mGy | xPN CNS | 0PN (CNS gr.) | -6.260 | 28.759 | 0.8281 |
| 11 | 10 mGy | xPN CNS | 0PN (CNS gr.) | -10.882 | 28.759 | 0.7059 |
| 12 | 100 mGy | xPN CNS | 0PN (CNS gr.) | -0.622 | 28.759 | 0.9828 |

| **C** | **dose** | **contrast** |  | **estimate** | **s.e.m.** | **p value** |
| --- | --- | --- | --- | --- | --- | --- |
| 1 | 2.5 mGy | xPN (S4B) | 0PN (S4B) | -34.961 | 8.418 | 3.47E-05 |
| 2 | 5 mGy | xPN (S4B) | 0PN (S4B) | -2.367 | 8.431 | 0.7790 |
| 3 | 10 mGy | xPN (S4B) | 0PN (S4B) | -0.259 | 8.694 | 0.9762 |
| 4 | 100 mGy | xPN (S4B) | 0PN (S4B) | 0.024 | 8.163 | 0.9976 |
| 5 | 2.5 mGy | xPN (S4C) | 0PN (S4C) | -34.977 | 9.687 | 0.0003 |
| 6 | 5 mGy | xPN (S4C) | 0PN (S4C) | -5.702 | 9.551 | 0.5506 |
| 7 | 10 mGy | xPN (S4C) | 0PN (S4C) | -1.738 | 9.773 | 0.8589 |
| 8 | 100 mGy | xPN (S4C) | 0PN (S4C) | 0.432 | 9.616 | 0.9642 |

## Declarations

### Ethics approval

The study was approved by the Ethics Committee of the Medical Association of Rhineland-Palatinate (no. 837.440.03 (4102) and no. 837.262.12 (8363-F)). All applicable institutional and governmental regulations concerning the ethical use of human volunteers were followed. Written informed consent to use primary fibroblasts for research purposes was obtained from all participants.

### Availability of data and materials

Supplemental Information accompanies this paper. The datasets of this article are either included within the article and its additional files, or are available upon request. Data sharing restrictions apply to the availability of the German Childhood Cancer Registry and KiKme study data described in this paper.

The ImageJ tool Cellect used for image processing and the software AutoFoci to automatically count foci in single cell images (including all relevant settings for this study) are described in Lengert *et al.* [21] and are freely available at https://github.com/nleng/AutoFoci.

### Author contributions

M.L. conceived the project, R.N.C. and C.S. performed experiments; J.M., R.N.C., C.S., A.S. and M.L. analyzed data; D.G., S.Z., P.S.-K., T.H., M.M., M.B. and H.S were involved in the selection of the KiKme study participants and recruitment; J.M. and M.L. designed and wrote the paper. All authors contributed to the interpretation of findings and the final manuscript.

## Acknowledgments

We thank the German Childhood Cancer Registry (Claudia Spix and Desiree Grabow), dermatologists and physicians of the University Medical Centre of the Johannes Gutenberg University Mainz, Germany and dermatologists at other locations throughout Germany involved in taking skin biopsies.

This work was supported by the Federal Ministry of Research, Technology and Space (02NUK016A-D & 02NUK042A-D).

---

**Figure S1:**
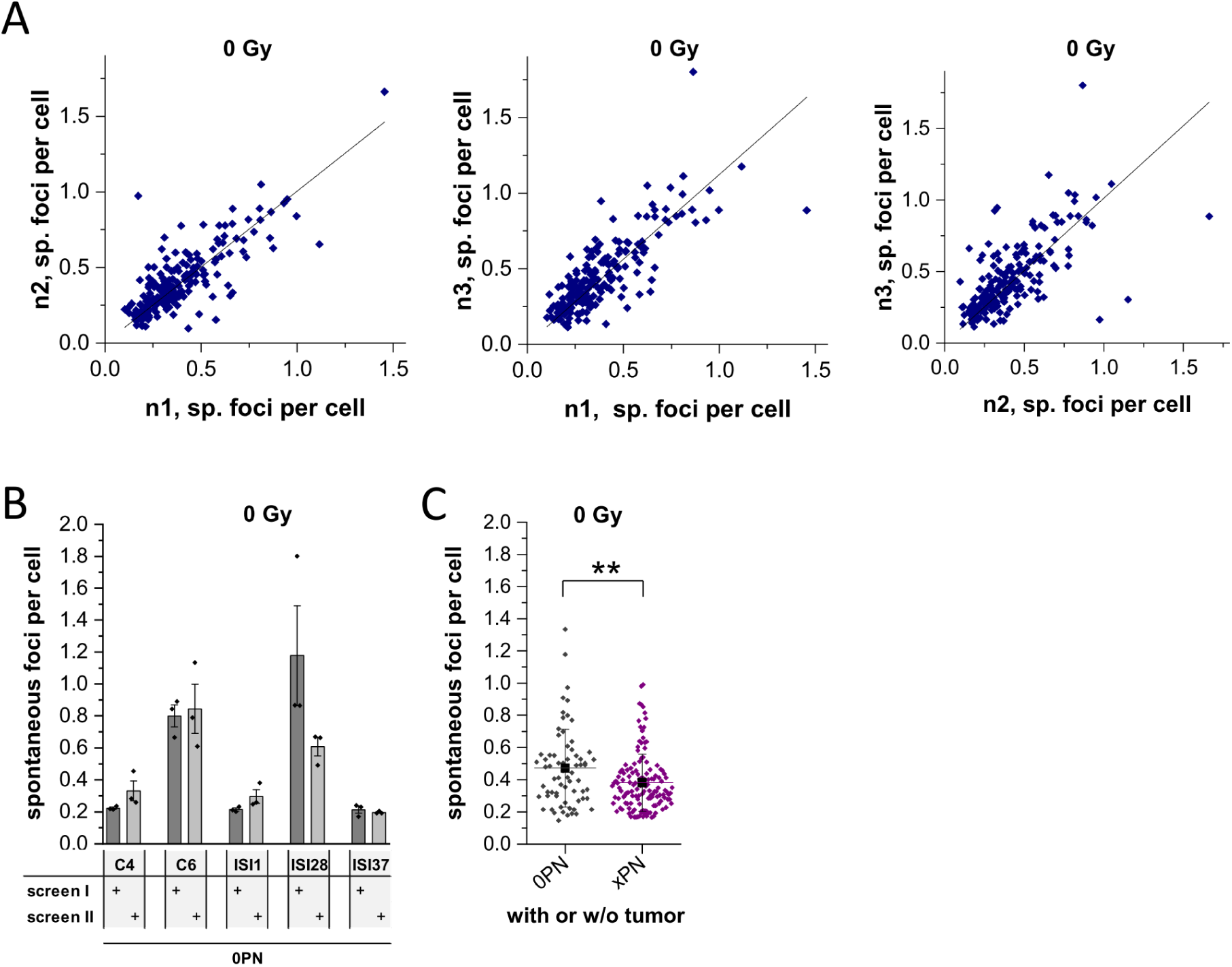
Accuracy and reproducibility of screening data. (**A**) Inter-experimental analysis of the data in figure 1C using a linear regression and a Spearman rank test. Each point represents the result of one individual from a single experiment. The slope of the curve, the quality criterion R^2^ and the Spearman rank coefficient (SRC) are: n1 vs. n2: f(x)= 1.00x (±0.022), R^2^ = 0.91, SRC = 0.75; n1 vs. n3: f(x)= 1.12x (±0.025), R^2^ = 0.91, SRC = 0.79; n2 vs. n3: f(x)= 1.01x (±0.029), R^2^ = 0.86, SRC = 0.71. (**B**) Validation of the data using a second screen with 5 samples. Data for individual experiments, the mean and s.e.m. are shown. Data for screen I are taken from figure 1C. (**C**) Analysis of spontaneous foci separated for 68 cancer-free individuals and 136 childhood cancer survivors. Each point represents one individual. The population mean and s.d. are shown in black. Statistical analysis was performed using a linear mixed model, and all statistical parameters are described in table 2A. 0PN, cancer-free individuals; xPN, individuals surviving at least one primary neoplasm in childhood; **, p ≤ 0.01.

**Figure S2:**
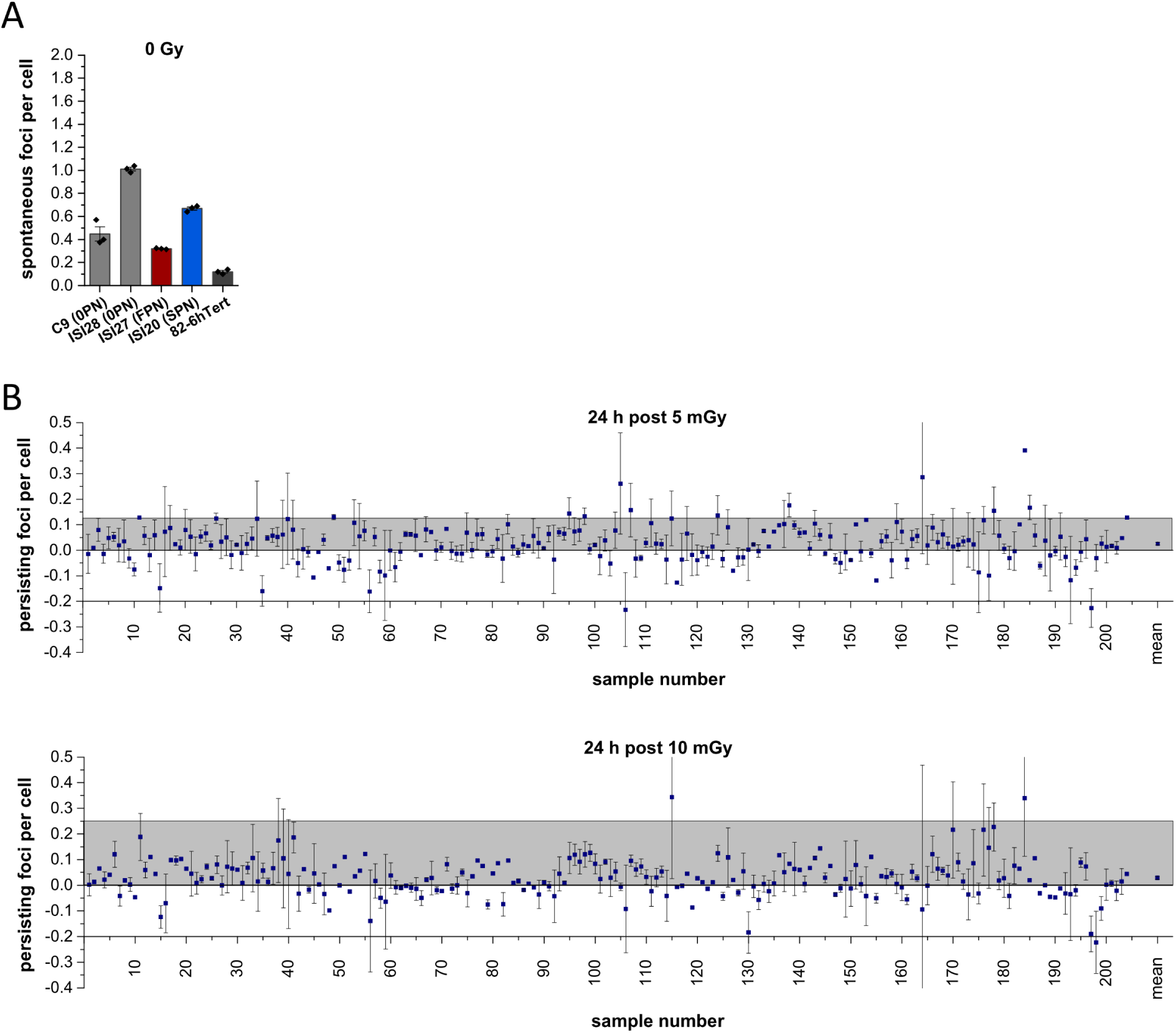
Spontaneous and persisting foci levels. (**A**) Spontaneous foci numbers in cells from the four selected individuals and an established cell line used for the analysis in figure 2C. Data for individual experiments, the mean and s.e.m. are shown. (**B**) Persisting foci after irradiation with 2.5 and 100 mGy. The data from panel 2B is replotted, showing the average number of persisting foci for the 204 individuals and the s.e.m. if three independent measurements are available. The population mean and s.e.m. are shown as the last point in each row. 0PN, cancer-free individuals; FPN, first primary neoplasm; SPN, second primary neoplasm.

**Figure S3:**
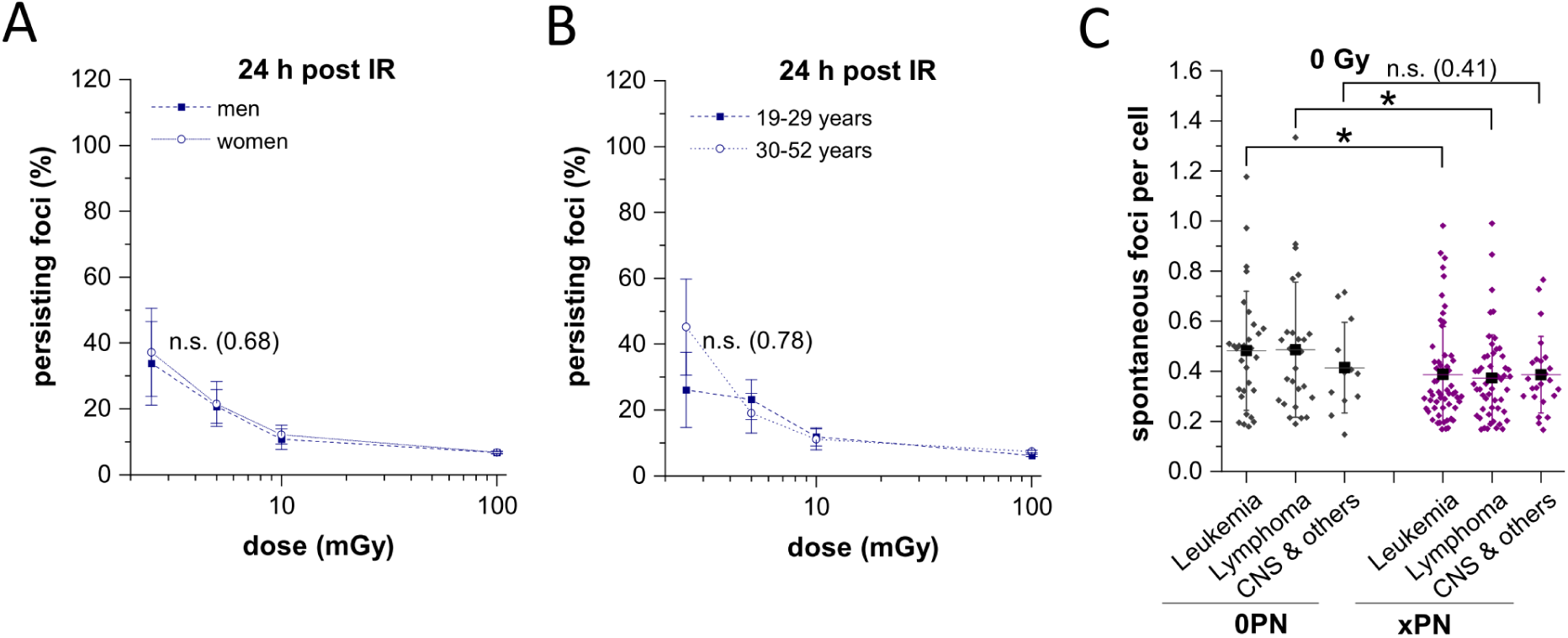
DSB repair after low-dose radiation is independent of age and sex. (**A, B**) Fraction of persisting foci as a function of dose separated by sex, with 99 males and 105 females (A), or for 99 individuals between 19-29 years and 105 individuals between 30-52 years of age at recruitment (B). The population mean and s.e.m. are shown. (**C**) Analysis of spontaneous foci for 60 individuals with leukemia, 52 individuals with lymphoma and 24 individuals with ‘CNS and other tumor entities’ as the first primary neoplasm, together with the matched 30, 26 and 12 cancer-free individuals. Each point represents one individual. The population mean and s.d. are shown in black. Statistical analysis was performed using a linear mixed model, and all statistical parameters are described in tables 4A (for panels A and B) and 2B (for panel C). 0PN, cancer-free individuals; xPN, individuals surviving at least one primary neoplasm in childhood; *, p ≤ 0.05; n.s., not significant.

**Figure S4:**
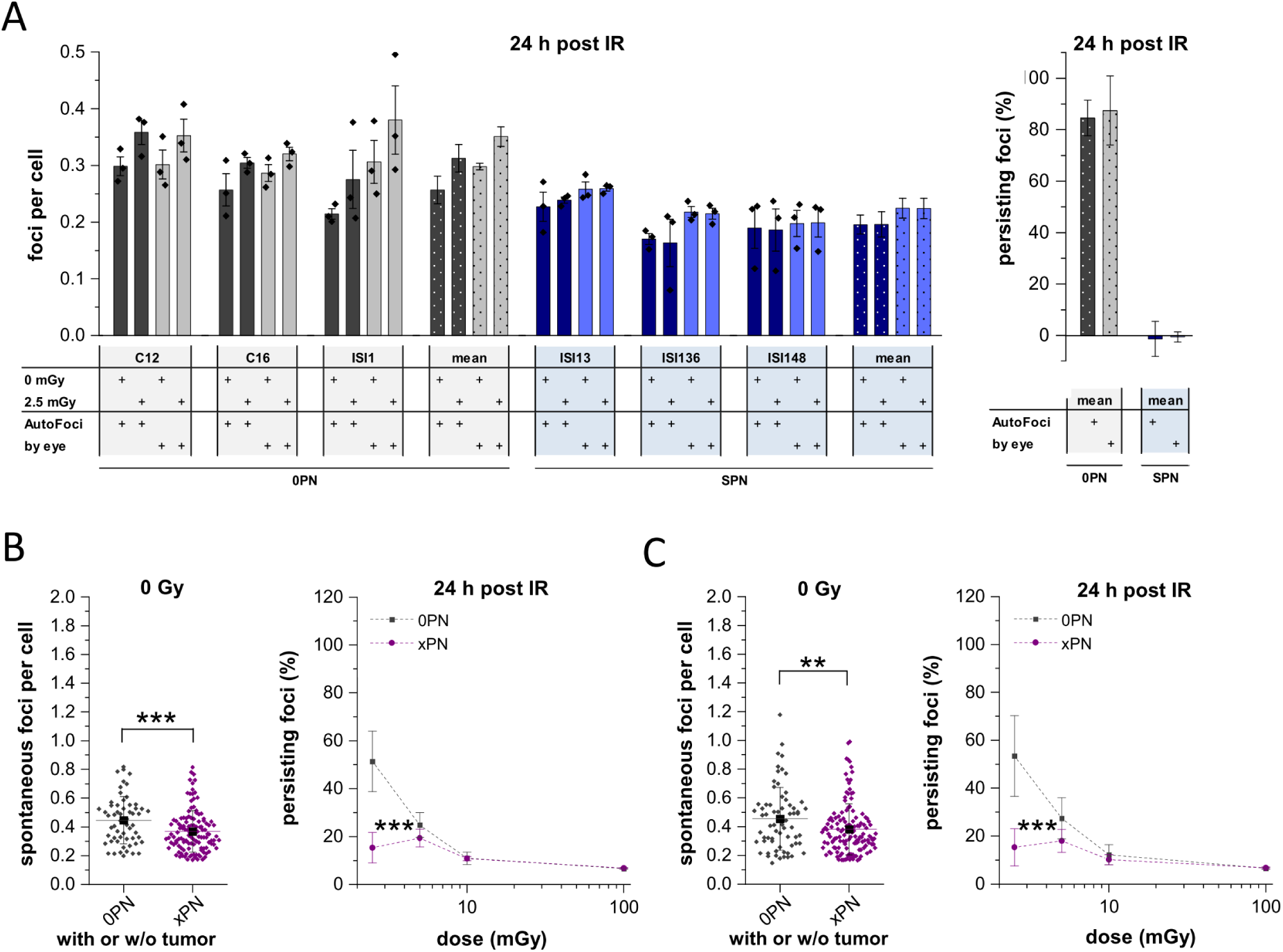
Verification of differences between the groups in DSB repair after low-dose radiation. (**A**) Verification of foci counting by AutoFoci and observed effects at 2.5 mGy. We selected three cancer-free individuals and three with a SPN, representing the differences in both spontaneous foci and repair efficiency observed within the entire analyzed group. Data for individual experiments, the mean and the s.e.m. are shown, together with the resulting population mean and the s.e.m. (left). The fraction of persisting foci was also calculated (right). The population mean and s.e.m. are shown. Data for ‘AutoFoci’ are taken from figures 1C and 2A. (**B**, **C**) Sensitivity analysis for the data in figures 1F and 3B. Either the 5 highest and 5 lowest values (B) or the values outside the 3x interquartile range (C) were deleted, for both groups and each dose separately. The population mean and s.d. (left panels in B and C) or s.e.m. (right panels in B and C) are shown. Statistical analysis was performed using a linear mixed model, and all statistical parameters are described in tables 2C and 4C. 0PN, cancer-free individuals; xPN, individuals surviving at least one primary neoplasm in childhood; SPN, second primary neoplasm; **, p ≤ 0.01; ***, p ≤ 0.001; n.s., not significant.

